# Olfactory sensory neurons mediate ultra-rapid antiviral immune responses in teleosts in a TrkA-dependent manner

**DOI:** 10.1101/464214

**Authors:** Ali Sepahi, Aurora Kraus, Christopher A Johnston, Jorge Galindo-Villegas, Cecelia Kelly, Diana García-Moreno, Pilar Muñoz, Victoriano Mulero, Mar Huertas, Irene Salinas

## Abstract

The nervous system is known to regulate host immune responses. However, the ability of neurons to detect danger and initiate immune responses at barrier tissues is unclear. Vertebrate olfactory sensory neurons (OSNs) are located in direct contact with the external environment and therefore directly exposed to pathogens. Here, we report that nasal delivery of rhadboviruses induced apoptosis in crypt OSNs in rainbow trout olfactory organ (OO) via the interaction of the OSN TrkA receptor with viral glycoprotein. This signal resulted in pro-inflammatory responses in the OO and dampened inflammation in the olfactory bulb (OB). CD8α^+^ cells infiltrated the OO within minutes of nasal viral delivery and this response was abrogated when TrkA was blocked. Infiltrating CD8α^+^ cells originated from the microvasculature surrounding the OB and not the periphery. Ablation of crypt neurons in zebrafish resulted in increased susceptibility to rhabdoviral challenge. Our results, therefore, indicate a novel function for OSNs as a first layer of pathogen detection in vertebrates.

## Introduction

The interactions between the nervous system and the immune system are multiple and complex. Both systems are specialized in sensing and responding to environmental signals and, evolutionary speaking, these two systems may have originated from a common ancestral precursor (Arendt, 2008). Sensory neurons have been shown to participate in immune responses in several animal models. For instance, in mice, nociceptor sensory neurons can be directly activated by bacteria to control pain and modulate inflammatory responses (Chiu et al., 2013; Pinho-Ribeiro et al., 2017) at several sites including the skin, joints, lungs and gastrointestinal tract (Pinho-Ribeiro et al., 2017). Moreover, sensory neurons are critical for suppressing innate immune responses triggered by pathogens and restore host homeostasis in invertebrates (Sun et al., 2012).

Vertebrate olfactory sensory neurons (OSNs) rapidly sense chemical stimuli present in the environment and transduce odorant-encoded signals into electrical signals that travel to the olfactory bulb (OB) via the olfactory nerve, where they are integrated and transferred to other parts of the central nervous system (CNS). OSNs are one of the few neurons in the vertebrate body that are in direct contact with the external environment, yet the interactions between microbes and OSNs remain unknown. OSNs are also in close proximity to a local network of immune cells known as the nasopharynx-associated lymphoid tissue (NALT) which is present in both teleosts and mammals (Sepahi and Salinas, 2016; Tacchi et al., 2014). The cross-talk between OSNs and NALT during the course of an immune response has not yet been investigated.

Teleost fish are known to have four different OSN types: ciliated, microvillous, crypt, and kappe neurons (Ahuja et al., 2014). The specific odors recognized by crypt neurons and their function are still enigmatic although recent evidence suggests that these neurons are responsible for kin recognition in zebrafish (Biechl et al., 2016). Crypt neurons only express one type of olfactory receptor, the vomeronasal receptor 1-like Ora4 and can be identified by their tropomyosin-related kinase A receptor (TrkA) immunoreactivity (Ahuja et al., 2013; Catania et al., 2003; Germana et al., 2004). The interaction between TrkA and endogenous ligands, such as nerve growth factor (NGF), induces internalization of TrkA into endosomes (Grimes et al., 1996). While TrkA activation by NGF regulates neuronal differentiation, growth and survival (Cattaneo and McKay, 1990; Sofroniew et al., 2001), previous studies have shown the ability of pathogens to hijack the TrkA system to infect hosts. For instance, *Trypanosoma cruzi* activates TrkA receptors and uses TrkA as a vehicle for host cell invasion (de Melo-Jorge and PereiraPerrin, 2007). Additionally, herpes simplex virus (HSV2) G protein modulates TrkA in order to guide axons to the infection site and facilitate neuronal infection (Cabrera et al., 2015).

Many neurotropic viruses exploit the olfactory route to infect CNS tissues (Koyuncu et al., 2013; Mori et al., 2005). In this context, murine studies suggest that nasal infection with neurotropic viruses can stimulate immune responses in the olfactory organ (OO) as well as the OB within hours of virus delivery (Leyva-Grado et al., 2010; Majde et al., 2007). Thus, currently, the OB in mice is considered as an immune effector organ capable of eliciting pro-inflammatory immune responses and containing pathogen infections to protect the CNS (Durrant et al., 2016). These local immune responses are isolated from the systemic immune compartment since the blood brain barrier (BBB) remains intact early during viral infection (Bi et al., 1995; D’Agostino et al., 2012).

In this study, we report that crypt neurons expressing TrkA are fast sensors of viruses in the olfactory mucosa and critical regulators of antiviral immune responses in teleost fish. These sensory neurons enter apoptosis within minutes of nasal delivery of infectious hematopoietic necrosis virus (IHNV), an aquatic rhabdovirus with neurotropic characteristics (LaPatra et al., 1995), via the interaction of the TrkA receptor neuron with viral glycoprotein (G protein). Upon exposure to virus, TrkA-dependent neuronal activation rapidly elicits pro-inflammatory immune responses in the OO and dampens inflammation in the OB. Furthermore, viral sensing by crypt neurons induces infiltration of CD8α ^+^ T cells from the microvasculature surrounding the OB into the OO and specific ablation of crypt neurons results in increased susceptibility to rhabdoviral challenge. Our results reveal that pathogen sensing by sensory neurons precedes that of the immune system and that initial signals derived from neurons allow quick orchestration of immune responses between the nasal mucosa and the CNS.

## Results

### Nasal delivery of neurotropic virus induces caspase-3 dependent apoptosis fo TrkA^+^ crypt neurons

Neurotropic viruses such as rhabdoviruses can infect OSNs and transneuronally infect the CNS (Koyuncu et al., 2013; Mori et al., 2005). Viruses can also cause caspase-dependent apoptosis in neurons via NGF signals (Allsopp et al., 1998; Chou et al., 2000; Gomes-Leal et al., 2006; Lavoie et al., 2005). We first confirmed that anti-TrkA antibody exclusively labels crypt neurons in trout. As previously reported, TrkA^+^ cells in the OO of trout had typical morphology and apical localization of crypt cells (Figure 1A). Immunoblotting of total tissue lysates with rabbit anti-TrkA antibody showed a band at the expected size of ~140kDa in OO and brain but not in the head-kidney (HK), the main hematopoietic tissue in bony fish (Figure S1A). Microscopy results confirmed the absence of TrkA^+^ cells in HK (Figure S1B) indicating trout immune cells are not TrkA^+^. When we delivered live attenuated IHNV intranasally (IN) into rainbow trout, we observed a significant decrease in the number of TrkA^+^ crypt cells in the OO compared to control fish 15 min, 1 h and 1 day after treatment (Figure 1 A, B, E & L). The number of TrkA^+^ crypt cells returned to basal levels by day 4 suggesting that that replacement from progenitors takes approximately 4 days to complete. Loss of TrkA reactivity 15 min after viral delivery was associated with presence of apoptotic morphology in the remaining TrkA^+^ crypt cells (Figure 1C, G & I) compared to controls (Figure 1D, H & J). Moreover, staining with anti-caspase 3 antibody confirmed that crypt neurons were undergoing apoptosis through caspase 3 pathway at this time point (Figure 1F). Since TrkA signaling may result in cell death in sensory neurons (Nikoletopoulou et al., 2010), we pharmacologically blocked TrkA with the drug AG879. Nasal delivery of AG879 30 min prior to IHNV nasal delivery (Figure 1K) rescued 50% of TrkA reactivity in crypt neurons (Figure 1M). Combined, these results indicate that aquatic rhabdoviruses result in crypt neuron cells death in the OO of rainbow trout in a TrkA-dependent manner.

**Figure 1:**
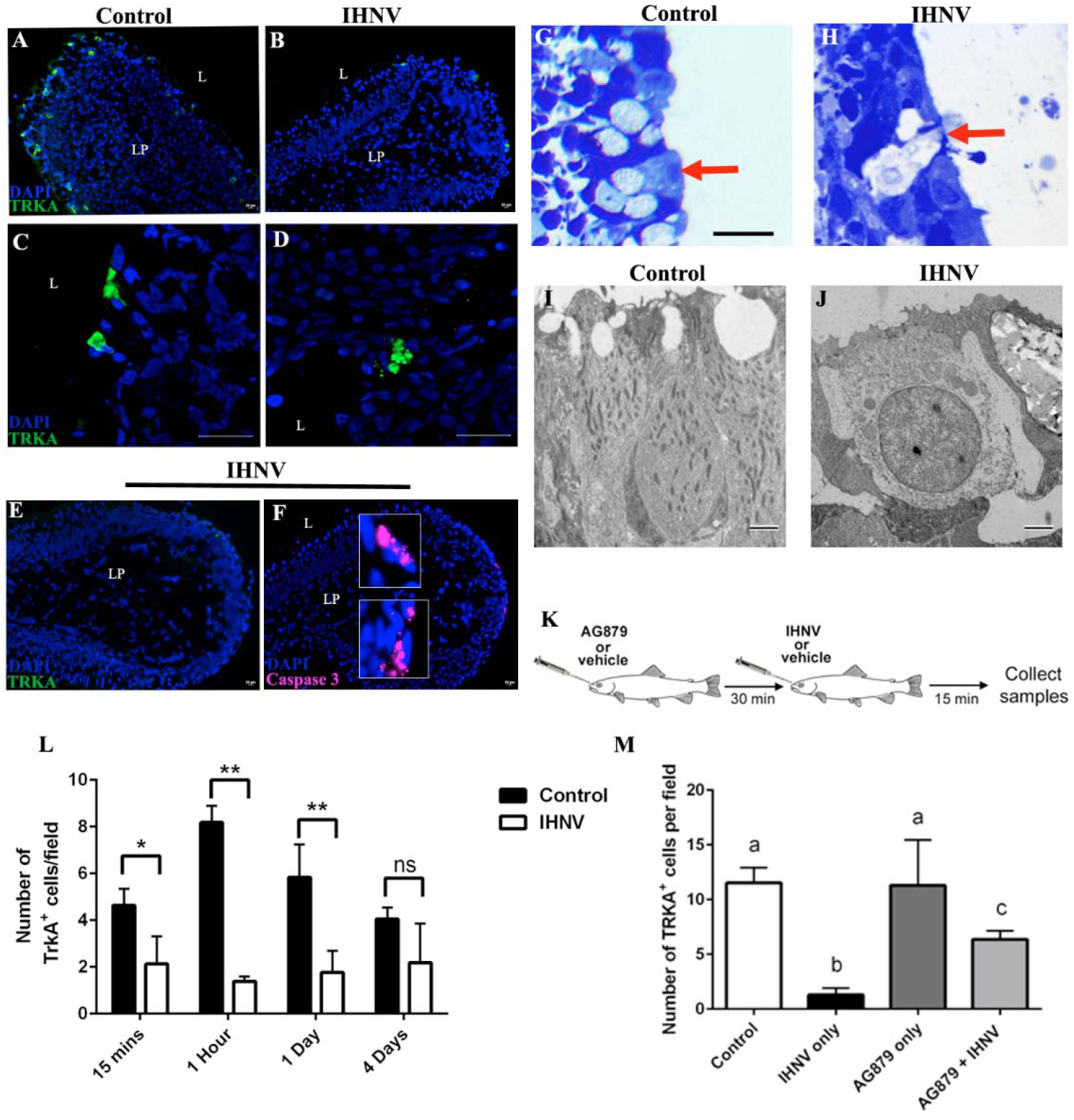
Nasal delivery of neurotropic virus induces caspase-3 dependent apoptosis to TrkA^+^ crypt neurons Immunofluorescence microscopy image of rainbow trout olfactory organ cryosections **(A)** Control and **(B)** 15 min after nasal IHNV delivery stained with anti-TrkA (FITC, green) show a decrease in the number of TrkA^+^ crypt neurons following IN delivery of IHNV. Nuclei were stained with nuclear stain DAPI (blue). Confocal microscopy images of rainbow trout olfactory organ cryosections **(C)** Control and **(D)** 15 min after nasal IHNV delivery stained with anti-TrkA (FITC, green) showing changes in TrkA reactivity with the characteristic morphology of cell apoptosis in TrkA^+^ cells following IN delivery of IHNV. Cell nuclei were stained with DAPI (blue). Scale bar, 20 μm. Immunofluorescence staining of an IHNV-vaccinated rainbow trout olfactory organ with anti-TrkA (FITC, green) **(E)** and anti-caspase 3 (Cy5, magenta) **(F)** showing co-localization of caspase-3 staining in low-TrkA^+^ crypt neurons. Scale bar, 20 μm. Cell nuclei were stained with DAPI DNA stain (blue). Results are representative of three independent experiments (N = 5) L, lumen; LP, lamina propria. Semithin sections of control (**G**) and nasal IHNV treated **(H)** rainbow trout olfactory organ indicate that crypt neurons undergo cell death within 15 min of viral delivery. Scale bar A-H, 20 μm. Transmission electron micrograph showing a crypt neuron in control trout olfactory epithelium **(I)** and a crypt neuron undergoing cell death in nasal IHNV treated rainbow trout 15 min after nasal vaccination **(J)**. Scale bar, 2 μm. **(K)** Schematic diagram of the experimental design used in TrkA-blocking experiment. AG879 (TrkA blocker) or vehicle were delivered IN and 30 min later IHNV or PBS (control) were delivered to the nasal cavity of trout. OO and OB were collected 15 min after IHNV delivery. **(L)** Quantification of the mean number of TrkA^+^ crypt neurons in control and nasal IHNV treated rainbow trout olfactory organ 15 min, 1 h, 1 day and 4 days after nasal viral delivery showing that TrkA reactivity begins to recover on day 4 as measured by immunofluorescence microscopy (N = 3). Results are expressed as mean ± SEM. Unpaired *t*-test *p < 0.05, **p < 0.01. **(M)** AG879 pre-treatment partially abolishes loss of IHNV-induced TrkA reactivity in crypt neurons (N = 3). Results are expressed as mean ± SEM and statistical. One-way ANOVA and a Tukey post hoc analysis test were performed to identify statistically significant differences among groups. P < 0.05.

### Rainbow trout smell neurotropic virus

Exposure of either live attenuated IHNV or culture medium used to grow the virus elicited strong olfactory responses and followed a dose-dependent pattern characteristic of activation of olfactory receptors (Figure 2A). Both the virus and the culture medium elicited highly sensitive olfactory responses, which could be detected up to a 1:10^5^ dilution by electro-olfactogram (EOG). However, IHNV elicited greater olfactory responses than medium at the 1: 100 dilution. Differences in the slopes of the linear dose responses also suggested activation of a different receptor set for each stimulus. Thus, we performed cross adaptation experiments in which the OO was continuously saturated with IHNV (adapted stimulus), and then measured olfactory responses to IHNV (self-adapted control) or a mix of IHNV with medium by EOG (see material and methods for details). We repeated the same experiment saturating the OO with medium and then measuring responses to medium alone or the IHNV and medium mix. If IHNV and medium activate different receptors, we would expect that OO saturated with IHNV will have a smaller olfactory response to a concentrated solution of IHNV due to fewer IHNV receptors available for activation. In turn, we would expect a greater response for the mix of IHNV and medium since medium-specific receptors but no IHNV-specific receptors would be available for activation. In agreement, cross-adaptation odorant assays showed that, after saturation of olfactory receptors with the adapting solution, normalized self-adapted controls had significant lower responses (26– 40 %) than the mixture of live attenuated IHNV and culture medium (Figure 2B), which implied different activation of receptors by the virus and medium, respectively.

**Figure 2:**
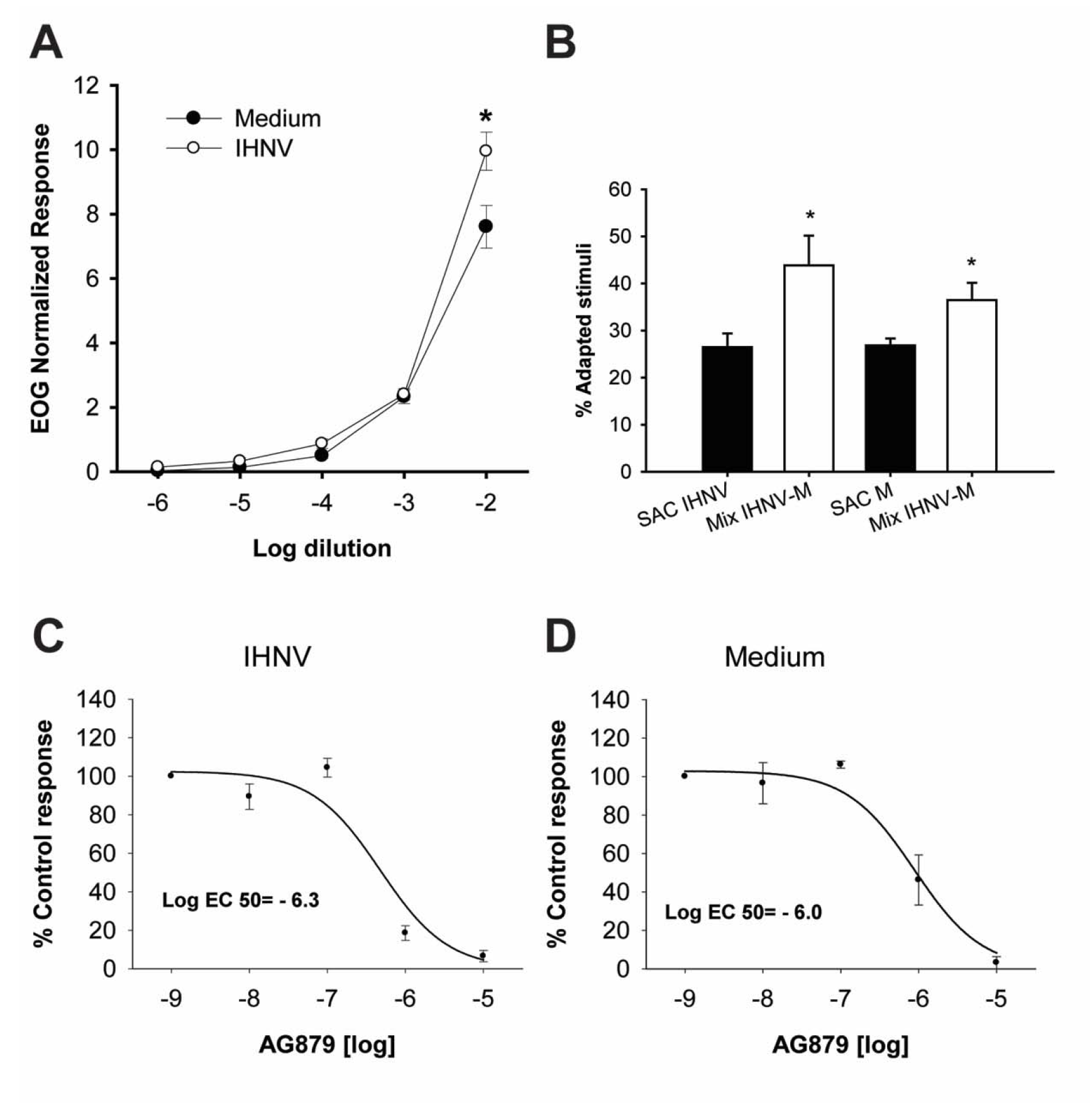
Rainbow trout smell neurotropic virus (**A**) Olfactory responses to live attenuated IHNV and medium where the virus was grown (negative control) produce different dose-response curves in rainbow trout measured by electro-olfactogram (EOG). Responses were normalized to the L-Serine control. Data are represented as the mean ± SEM (N = 8). Paired t-test showed significant differences at dilution 1: 100 (P < 0.05). (**B**) Live attenuated IHNV activates a set of receptors different than those activated by virus-free medium (negative control). Cross-adaptation experiments compared olfactory responses to live attenuated IHNV (self-adapted control, SAC) or a mix of live attenuated virus and the virus-free medium (Mix IHNV-M) when the olfactory epithelium was saturated with live attenuated IHNV (adapted stimuli) odors. Same experiments were performed using the IHNV culture medium as adapted stimuli. Paired t-test showed significant differences (p < 0.05) between both SAC and Mix. (**C**) AG879 treatment results in stronger inhibition of olfactory responses in live attenuated IHNV than in virus-free supernatant. (**D**) Total pharmacological inhibition of olfactory responses was achieved at concentrations of the drug >10^−5^ M. Paired t-test showed significant differences (p < 0.05) between EC_50_.

Since we hypothesized that viral detection is TrkA receptor-mediated, we expected a decrease of olfactory responses after nasal exposure to TrkA inhibitor AG879. Inhibition curves showed that AG879 affected the olfactory responses to virus and culture medium in concentrations of the drug as low as 10^−8^ M, with a total inhibition of activity at 10^−5^ M (Figure 2C & D). The inhibition of olfactory responses by the drug was stronger for the virus than the medium, with an inhibition of 50% of olfactory responses (EC50) by AG879 of 10^−6.3^ M, and 10^−6^ M for virus and medium respectively. Inhibitory curves also showed hormesis at 10^−7^ M for the virus but not for the medium, implying a positive effect of the drug for virus detection at very low concentrations. Combined, these experiments demonstrate the rainbow trout is able to smell viruses via TrkA signaling.

### Neurotropic viruses activate sensory neurons in the OO and OB in a TrkA-dependent manner

Studies in fish have demonstrated that pERK staining and *c-fos* gene expression are suitable markers of neuronal activation upon odorant exposure in OO and CNS (Dieris et al., 2017; Lau et al., 2011). However, whether viruses activate neurons in the OO and OB has not been investigated to date. Incubation of OO single cell suspensions with IHNV *in vitro* showed a significant increase in pERK labeling after 15 min as measured by flow cytometry (Figure S2A & B). These results were confirmed *in vivo* since we detected pERK staining in the OO and OB of IHNV-treated fish but not controls (Figure 3A-D). OB neuronal activation was due to viral-derived signals present in the OO since IHNV could not be detected in the OB 15 min after nasal delivery (Figure S3A-D). Interestingly, pERK positive cells in the OO of IHNV–treated fish were localized in the middle of the neuroepithelium and did not have a crypt neuron morphology, indicating that OSNs other than crypt neurons become activated following nasal viral delivery (Figure 3C). We also found a significant upregulation in *c-fos* expression in the OO but not the OB, in fish that received IHNV compared to controls (Figure 3E-G). Inhibition of TrkA signaling pathway with AG879 before IN IHNV delivery blocked neuronal activation as evidenced by the lack of pERK staining in the OO and OB and absence of *c-fos* up-regulation in the OO (Figure 3E-G). Combined, these data indicate that IHNV induces OSN activation through a TrkA-dependent pathway or, alternatively, that crypt neuron cell death results in activation of other OSNs.

**Figure 3:**
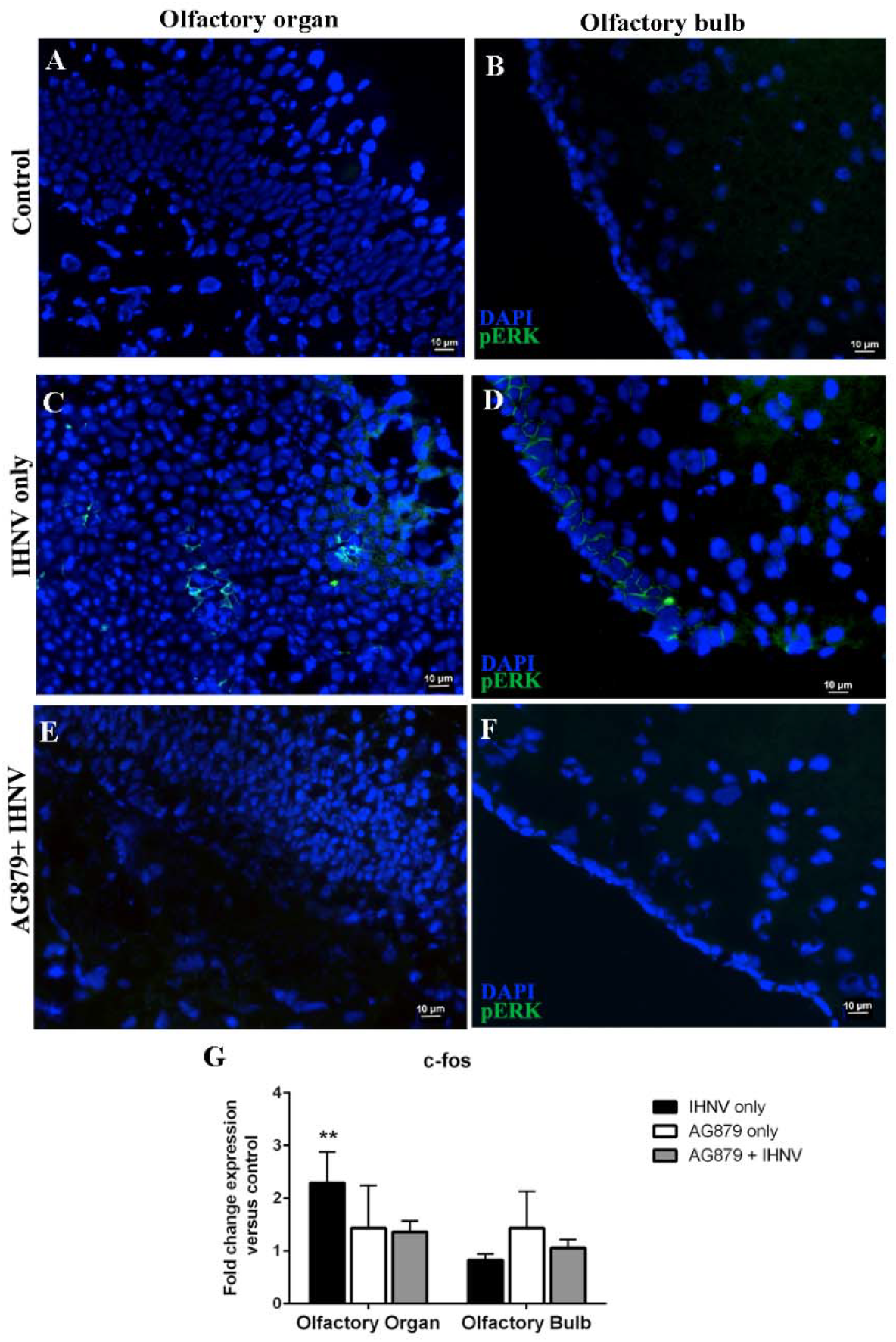
Nasal IHNV delivery activates sensory neurons in the OO and OB in a TrkA-dependent manner. Immunofluorescence staining of control rainbow trout OO (A) and OB (B) cryosections stained with anti-pERK (FITC, green) showing absence of neuronal activation. Immunofluorescence staining of OO (**C**) and OB (**D**) cryosections of trout nasally treated with IHNV stained with anti-pERK antibody (FITC, green) showing neuronal activation 15 min after viral delivery. Immunofluorescence staining rainbow trout OO (**E**) and OB (**F**) treated with AG879 + IHNV with anti-pERK showing AG879 inhibition of viral-induced neuronal activation in both OO and OB. Scale bar, 10 μm. For (A-F), cell nuclei were stained with DAPI DNA stain. (**G**) Gene expression levels of *c-fos* (neuronal activation marker) in control, nasal IHNV treated, AG879 only treated and IHNV + AG879 groups as measured by RT-qPCR. Gene expression levels were normalized to the housekeeping gene EF-1a and expressed as the fold-change compared to the control group using the Pfaffl method. Results are representative of three independent experiments (N = 5). Results were analyzed by unpaired t-test *p < 0.05, **p < 0.01.

### Nasal delivery of viruses results in ultra-rapid innate immune responses in the olfactory mucosa and the CNS in a TrkA-dependent manner

We previously reported that nasal delivery of live attenuated IHNV results in the recruitment of myeloid and lymphoid cells to the local nasal environment 4 days after treatment in trout (Sepahi et al., 2017; Tacchi et al., 2014). Here, we report that leukocyte recruitment occurs as early as 15 min after IHNV delivery as visualized by the enlarged lamina propria (LP) of the olfactory lamellae of IHNV-treated fish compared to control fish (Figure S4A-C). Histological changes in the OO were paralleled by changes in innate immune gene expression. Specifically, we observed a significant upregulation of *ck10*, a CCL19-like chemokine in rainbow trout (Sepahi et al., 2017), and *ptgs2b* in OO 15 min after IHNV delivery. In the OB, in turn, we observed a significant downregulation in expression of *ck10* and no significant change in expression of *ptgs2b* (Figure 4A). *ifng* expression was down-regulated both in the OO (2-fold) and the OB (4-fold) (Figure 4A) whereas no significant changes in *tnfa* expression were recorded in any tissue for any of the treatments. Importantly, when we pharmacologically blocked TrkA, we could revert IHNV-elicited changes in innate immune gene expression in both the OO and OB (Figure 4A). These results indicate antiviral pro-inflammatory immune responses in the nasal mucosa are accompanied by dampened antiviral immune responses in the OB and that both types of responses require TrkA activation in crypt neurons.

**Figure 4:**
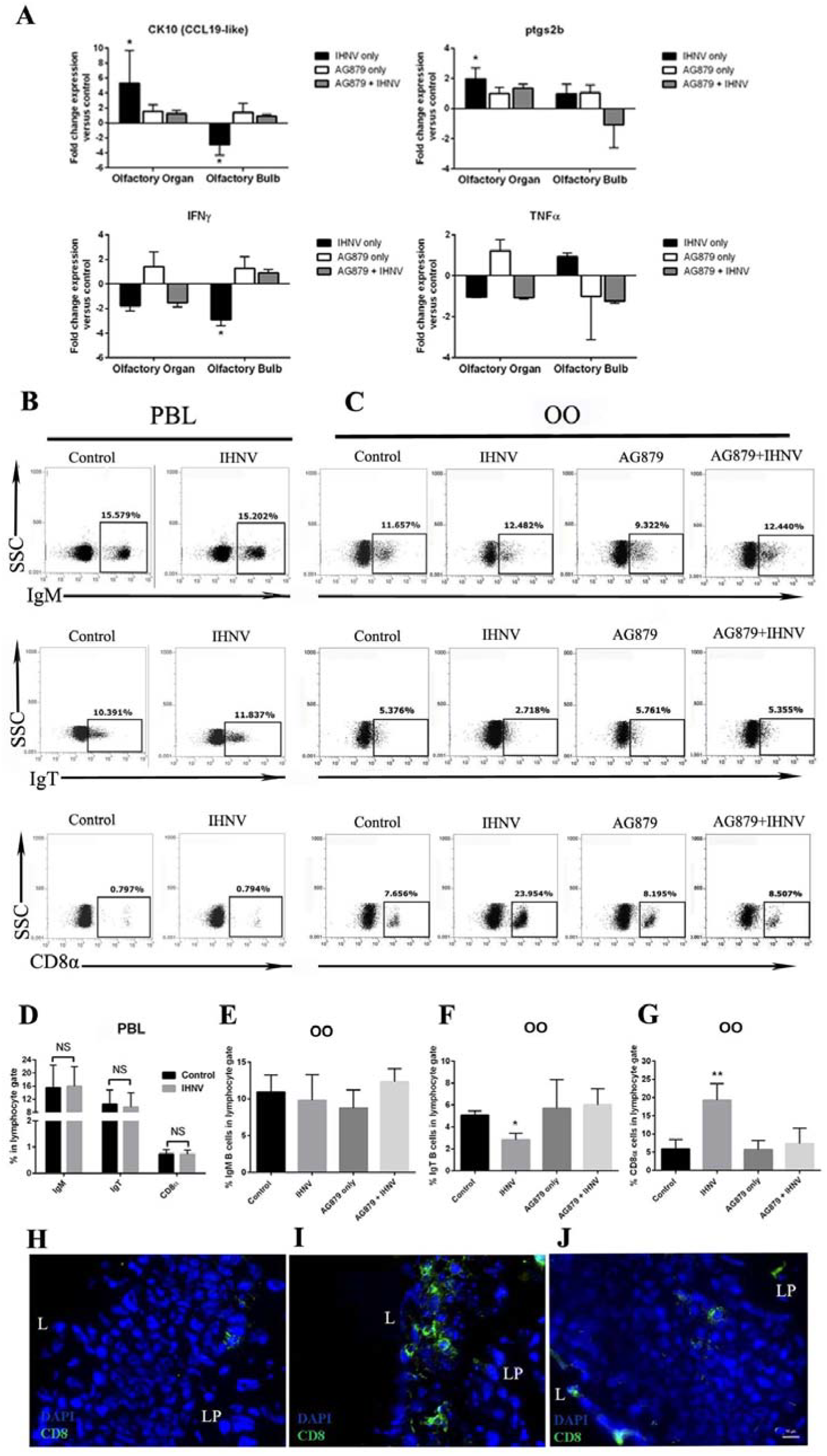
Viral nasal delivery results in ultra-rapid immune responses in the OO and OB of rainbow trout in a TrkA-dependent manner. **(A)** Gene expression levels of the chemokine *ck10* (CCL19-like), *ptgs2b*, *ifng* and *tnfa* in OO and OB in of control, IHNV-treated, AG879 treated and AG879 + IHNV treated trout. Gene expression levels were measured by RT-qPCR and normalized to the housekeeping gene *ef1a*. Data are expressed as the mean fold change compared to the control group using the Pfaffl method. Results are representative of three different experiments (N = 5). Results were analyzed by unpaired t-test *p < 0.05. **(B)** Representative dot plots of control and IHNV-treated rainbow trout PBLs stained with mouse anti-trout IgM, mouse anti-trout IgT and rat anti-trout CD8α showing the mean percentage of positive cells from the lymphocyte gate. **(C)** Representative dot plots of control and IHNV, AG879 and AG879 + IHNV trout OO lymphocytes stained with mouse anti-trout IgM, mouse anti-trout IgT and rat anti-trout CD8α showing the mean percentage of positive cells from the lymphocyte gate. **(D)** Quantification of flow cytometry data in **(B)** indicating no significant changes in the percentage of IgM^+^, IgT^+^ and CD8α^+^ cells in PBLs 15 min after nasal IHNV delivery. Results are representative of three independent experiments (N = 5). NS = not significant. **(E-G)** Quantification of flow cytometry data shown in **(C)** indicating significant decrease in the percentage of IgT^+^ B cells and increase in the percentage of CD8α+ T cells in the OO 15 min after nasal IHNV delivery and inhibition of such responses when AG879 is administered 30 min before IHNV treatment. Results are representative of three different experiments (N = 5). One-way ANOVA and a Tukey post hoc analysis test were performed to identify statistically significant differences among groups. *p < 0.05, **p < 0.01. **(H-J)** Immunofluorescence staining of a control **(H)** IHNV only **(I)** and AG879+IHNV **(J)** Rainbow trout OO cryosection stained with anti-CD8α (FITC, green). Scale bar, 10 μm. Cell nuclei were stained with DAPI DNA stain. L, lumen; LP, lamina propria. Results are representative of three different experiments (N = 5). Results were analyzed by unpaired t-test *p < 0.05.

### CD8α T cells rapidly infiltrate the olfactory organ in a TrkA-dependent manner

Gene expression data indicated that local immune responses in the OO of fish are initiated within minutes of viral exposure. We next evaluated changes in leukocyte populations in the OO and systemic circulation by flow cytometry following IHNV IN delivery. The percentages of trout IgM^+^, IgT^+^ B cells and CD8α^+^ T cells remained unchanged in in PBLs 15 min after nasal viral delivery compared to control fish (Figure 4B & D). In turn, the percentage of CD8α^+^ T cells increased from 7% in controls to 24% in IHNV-treated fish in the OO. Additionally, whereas no significant changes in IgM^+^ B cells were observed, we recorded a decrease in the percentage of IgT^+^ B cells after nasal viral exposure (Figure 4C, E, F & G). To test whether signals derived from crypt neurons are responsible for the observed infiltration of CD8α^+^ T cells into the OO, we blocked TrkA with AG879, as explained in Figure 1K. We found that AG879 treatment abolished both the decrease in the percentage of IgT^+^ B cells and the increase in CD8α^+^ T cells (Figure 4F & G). These results indicate that crypt neurons trigger ultra-rapid cellular immune responses against rhabodviruses in a TrkA-dependent manner.

### CD8α T cells infiltrates originate in the OB microvasculature

Neurotropic virus can hijack TrkA receptor to infect neurons and induce apoptosis (Allsopp et al., 1998; Chou et al., 2000; Gluska et al., 2014; Gomes-Leal et al., 2006; Lavoie et al., 2005). TrkA expressing crypt neurons project to a single target glomerulus in the OB (Ahuja et al., 2013) and we showed that nasal delivery of virus results in neuronal activation in the OO and OB. Moreover, we found that PBLs populations do not change following nasal viral delivery in trout. We therefore hypothesized that ultra-rapid infiltration of CD8α^+^ T cells in the OO might originate from a pool of lymphocytes present at the OB microvasculature that are recruited after neuronal signals. To test this, we collected the blood from the microvasculature surrounding the OB, 15 min after IN delivery of IHNV and evaluated changes in the percentage of cells within the lymphocyte gate as well as CD8α^+^ T cells, IgM^+^ and IgT^+^ B cells by flow cytometry. We observed a significant decrease in the percentage of cells within the lymphocyte gate 15 min after nasal delivery of IHNV (Figure 5A & C). Analyses of OB microvasculature leukocytes showed no significant changes in the percentage of IgM^+^ or IgT^+^ B cells but a significant decrease in the percentage of CD8α^+^ T cells from ~2% to ~0.5% (Figure 5B & D). Intravenous administration of FITC-conjugated dextran indicated that these effects occur without any changes in the blood barrier integrity of IHNV-treated fish (Figure S3E-H). Combined, these results indicate an ultra-rapid shunting of CD8α^+^ T cells from the OB to the OO.

**Figure 5:**
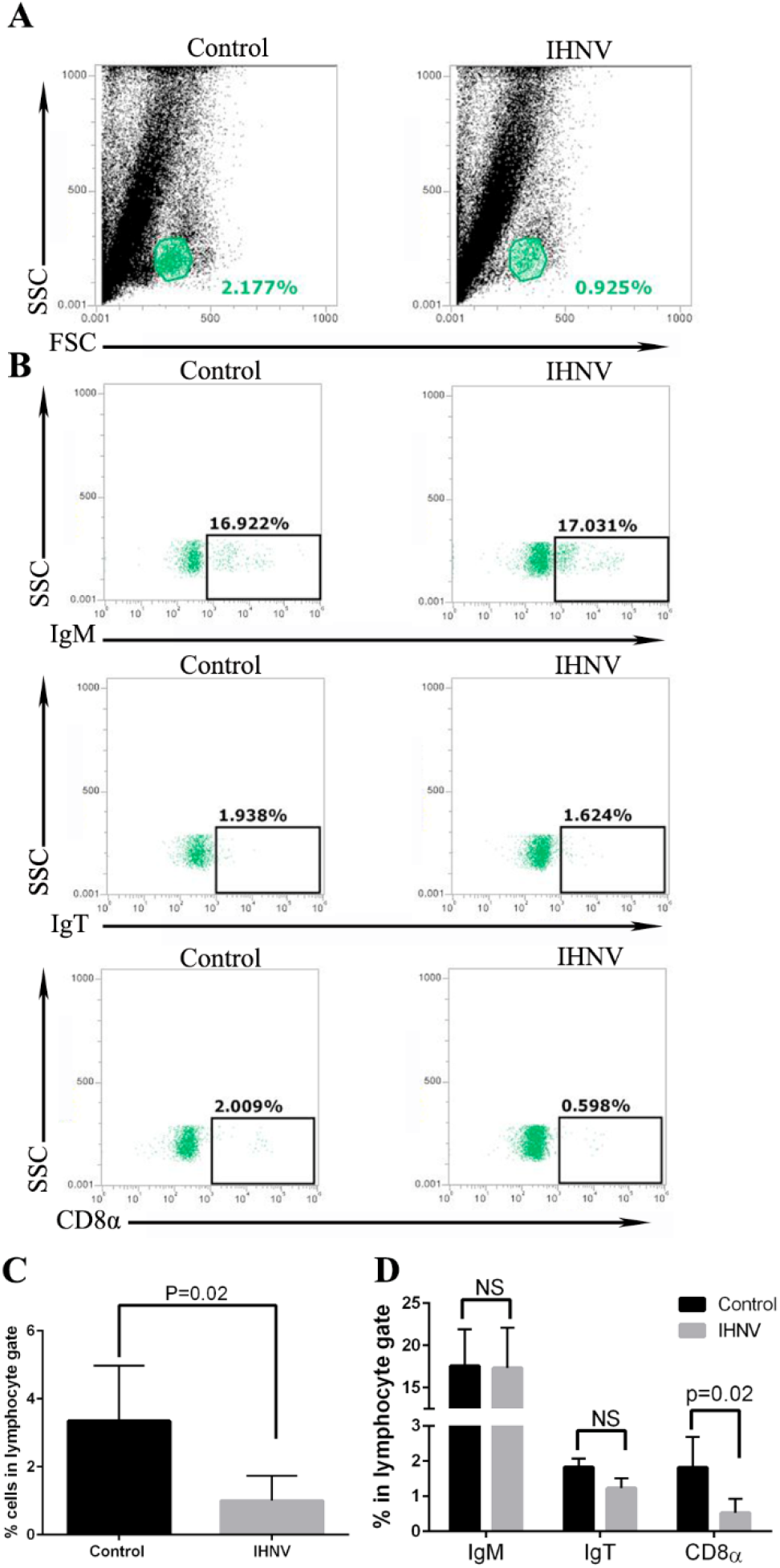
CD8α^+^ T cells infiltrating the trout OO originate from the OB microvasculature but not from peripheral blood. **(A)** Representative dot plots of control and IHNV-treated trout of cells obtained from the OB microvasculature showing the percentage of cells within the lymphocyte gate in each group. **(B)** Representative dot plots of cells obtained from the OB microvasculature of control and IHNV-treated rainbow trout stained with mouse anti-trout IgM, mouse anti-trout IgT and rat anti-trout CD8α showing the mean percentage of positive cells within the lymphocyte gate. **(C)** Quantification of flow cytometry data presented in **(A)** showing a significant decrease in the percentage of cells within the lymphocyte gate in IHNV-treated group compared to controls. **(D)** Quantification of flow cytometry data shown in **(B)** indicating a significant decrease in the percentage of CD8α+ cells in the OB microvasculature 15 min after nasal IHNV delivery. Results are representative of three different experiments (N = 5). Results were analyzed by unpaired t-test. NS = not significant.

### The interaction between IHNV viral glycoprotein (G protein) and TrkA is necessary for the onset of nasal antiviral immune responses

Since HSV Secreted G protein has been previously shown to interact with mouse TrkA (Cabrera et al., 2015), we hypothesized that IHNV G protein may be the ligand for TrkA in rainbow trout crypt neurons. Amino acid sequence analysis of vertebrate TrkA molecules showed a high (>50%) conservation among mouse, human and rainbow trout TrkA (Figure 6A) including amino acid sites known to interact with NGF (Wiesmann et al., 1999), whereas comparison of IHNV G protein and HSV secreted G protein indicated a low degree of amino acid conservation (Figure S5A). In order to test whether IHNV G protein alone is able to recapitulate IHNV-induced changes in crypt neurons and elicit nasal CD8α^+^ T cell immune responses, we produced FLAG-tagged IHNV G protein in a mammalian expression system (Figure S5B) and delivered 100 ng of recombinant IHNV G protein IN to rainbow trout. Microscopy results showed the co-localization of TrkA and IHNV G protein 15 min after administration of FLAG-tagged IHNV G protein (Figure 6B). We also observed a significant decrease in the number of TrkA^+^ cells 15 min after IN delivery of FLAG-tagged IHNV G protein (Figure 6C) similar to what we observed following IHNV treatment. Moreover, nasal delivery of FLAG-tagged IHNV G protein resulted in a significant increase in the number of CD8α^+^ T cells present in trout OO 15 min after nasal delivery (Figure 6D). To further confirm that IHNV G protein is responsible for TrkA-mediated nasal immune responses, we performed *in vivo* antibody neutralization experiments using a monoclonal anti-IHNV G protein antibody or a monoclonal anti-IHNV N protein antibody as a control. Blocking IHNV G protein but not N protein rescued the loss of TrkA reactivity in crypt neurons and abolished the infiltration of CD8α^+^ T cells into the trout OO 15 min after IN delivery (Figures 6E & F). These experiments demonstrated that the interaction between viral G protein and the crypt neuron TrkA receptor is necessary and sufficient to elicit OO immune responses.

**Figure 6:**
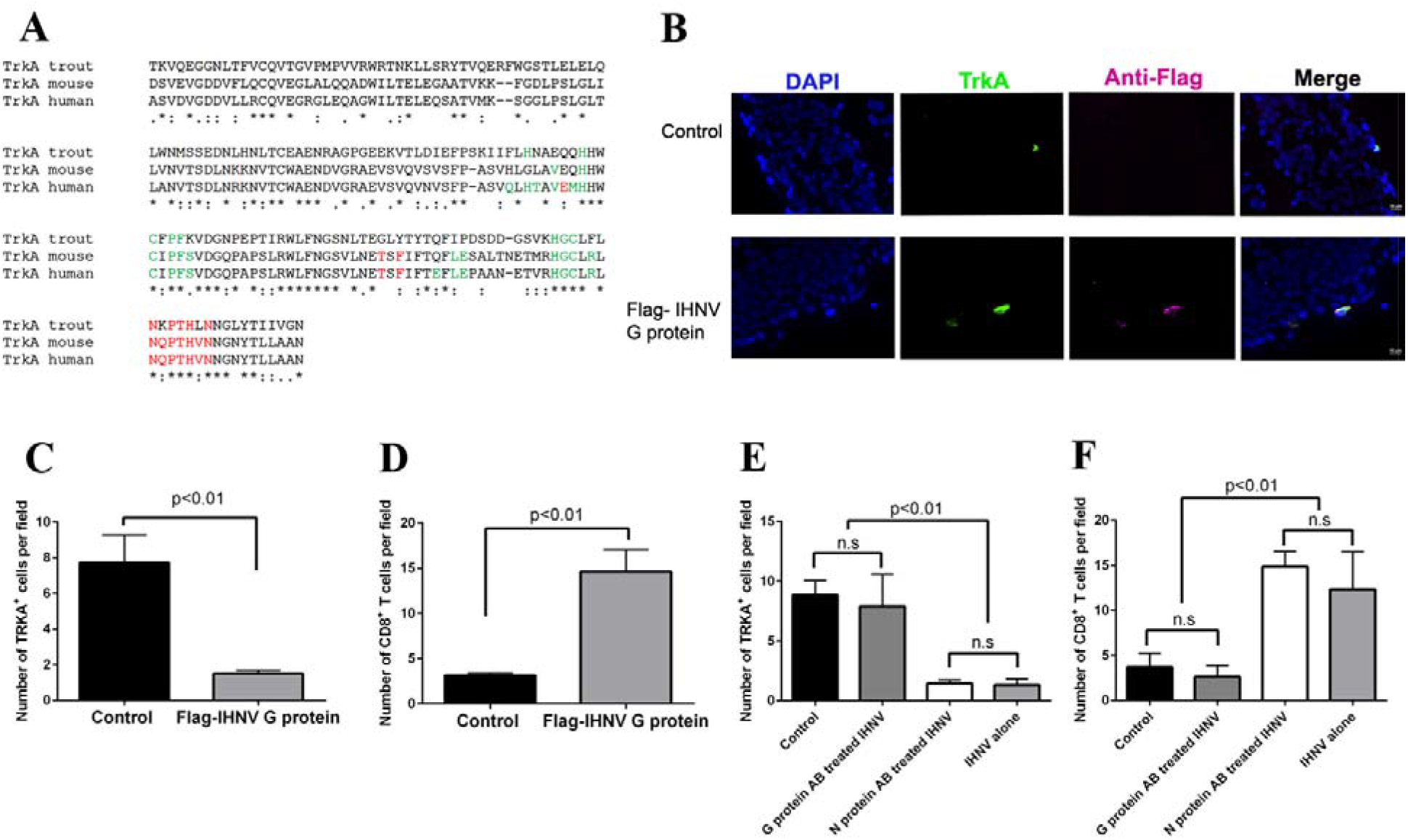
The interaction between viral glycoprotein (G protein) and crypt neuron TrkA i necessary for inducing crypt neuron-mediated nasal immune responses in trout. **(A)** Multiple sequence alignment (performed with CLUSTALW http://align.genome.jp/) of rainbow trout, mouse and human TrkA domain 5 (domain known to interact with cognate ligand) showing conservation of aa at sites previously described to be critical for NGF binding to TrkA. **(B)** Nasal delivery of recombinant IHNV G protein recapitulates IHNV-induced changes in crypt neurons and CD8 T cell immune responses. Immunofluorescence staining of trout olfactory organs 15 min after receiving PBS or 100 ng of recombinant FLAG-tagged IHNV G protein intranasally stained with anti-TrkA (FITC, green), anti-FLAG (Cy3, magenta) and DAPI (blue) showing the co-localization of TrkA and IHNV G protein in the FLAG-tagged IHNV G protein delivered group but not controls. **(C)** Quantification of the mean number of TrkA^+^ crypt neurons in the OO of control trout and trout that received recombinant FLAG-tagged IHNV G protein IN (N = 3). **(D)** Quantification of the mean number of CD8α^+^ T cells in control and FLAG-tagged IHNV G protein treated rainbow trout OO (N = 3) by immunofluorescence microscopy. (**E)** *In vivo* antibody blocking of IHNV G protein reverts IHNV-induced changes in crypt neurons and CD8 T cell immune responses. Live attenuated IHNV was incubated with anti-IHNV G protein monoclonal antibody, anti-IHNV N protein monoclonal antibody or not treated for 30 min at RT prior to *in vivo* nasal delivery. Quantification of the mean number of TrkA^+^ crypt neurons by immunofluorescence microscopy in control, anti-G protein antibody treated + IHNV, anti-N protein antibody treated IHNV and IHNV alone in the OO of rainbow trout (N = 3). **(F)** Quantification of the mean number of CD8α^+^ T cells by immunofluorescence microscopy in the OO of control, anti-G protein antibody treated + IHNV, anti-N protein antibody treated IHNV and IHNV-treated rainbow trout (N = 3). Results are representative of two independent experiments (N = 3). One-way ANOVA and a Tukey post hoc analysis test were performed to identify statistically significant differences among groups. *p < 0.05, **p < 0.01.

### Crypt neurons are involved in survival to rhabdoviral infection

Our results thus far provided evidence for the role of crypt neurons in the immune crosstalk between the olfactory organ and the CNS. Next, we asked whether viral detection by crypt neurons is necessary for survival against rhabdoviral infection using a a zebrafish model (Figure 7A). To that end, we generated a transgenic zebrafish line which expressed the Gal4.VP16 transactivator under the *ora4* promoter and the GFP under the heart specific promoter *cmlc2* for rapid screening under the fluorescence microscope (Figure 7A). Crossing of this line with the a line which expresses bacterial nitroreductase fused to mCherry revealed the presence of *ora*+ crypt neurons from 2 dpf onward (Figure 7A & 7B).. Addition of the prodrug metronidzole (Mtz) into the zebrafish water tanks at 2 dpf for 24 h resulted in 100% ablation of crypt neurons, that started to regenerate 5 days later (7 dpf) (Figure 7B & 7C). Infection of and ^ora4+^ crypt neuron ablated zebrafish with spring viraemia of carp virus (SVCV) resulted on defects in CCL19-like expression patterns in response to infection (Figure 7D), as measured by RT-qPCR. Crypt neuron ablation resulted in no significant differences in SVCV viral loads 15 min after exposure but increased SVCV loads 2 days post-infection (dpi) (Figure 7E). Importantly, challenge with SVCV revealed that, in the absence of crypt neurons, zebrafish are more susceptible to viral infection (Figure 7F). These results demonstrate that crypt neurons induce immune responses in response to viral infection and that these responses are essential for viral clearance and host survival.

**Figure 7:**
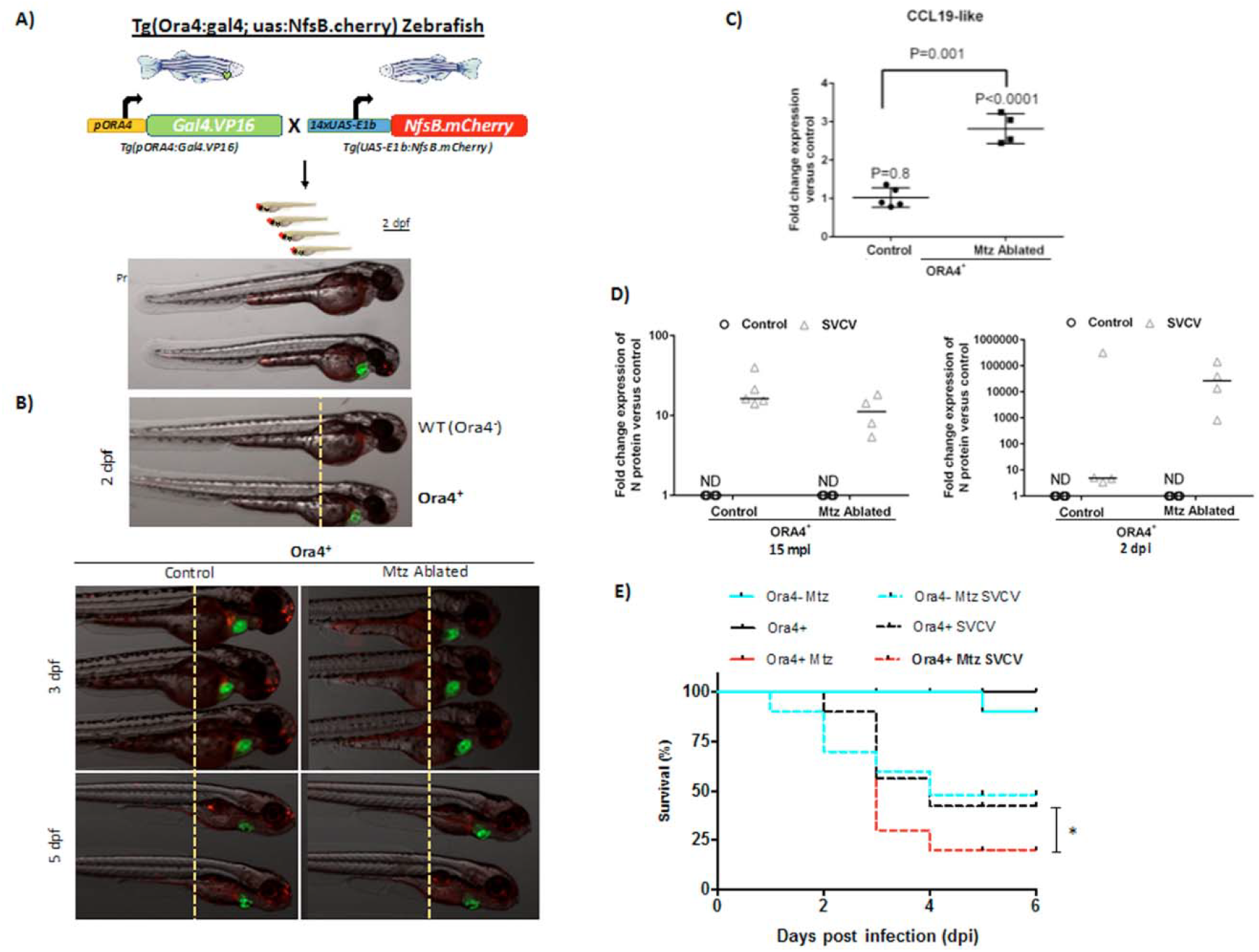
Ablation of crypt neurons results in increased susceptibility to rhabdoviral infection in zebrafish. (**A, B**) Schematic representation of the generation of *ora4* transgenic zebrafish and ablation of crypt neurons by delivery of the prodrug metronidazole (Mtz) into the water for 24 h. (**C)** Expression of CCL19-like in whole zebrafish larvae 15 min after infection with SVCV as measured by RT-qPCR. Each symbol represents a pool of 10 larvae. Results are expressed as the fold-change in expression compared to uninfected controls. Expression levels were normalized to the *rps11* as house-keeping gene. Data are expressed as the mean fold change compared to the control group using the Pfaffl method. Results are representative of three different experiments (N = 5). Results were analyzed by unpaired t-test. (**D**) Relative SVCV viral loads as measured by N protein expression levels 15 min and 2 days after SVCV infection in ORA4^+^ zebrafish and ORA4^+^ + Mtz (ablated) zebrafish. Each symbol represents a pool of 10 larvae. Results are expressed as mean level of N protein expression measured by RT-qPCR compared to uninfected controls. ND: non-detectable. P-values were obtained by unpaired t-test. (**E**) Percent survival of wildtype (ORA4-) and transgenic (ORA4+) zebrafish larvae that were ablated (+ Mtz) or not in response to SVCV infection. Results are representative of two independent experiments (N = 30 per group). Statistical analysis was performed by Gehan-Breslow-Wilcoxon method (p < 0.05).

## Discussion

The multifaceted, complex and bidirectional interactions between the nervous and the immune system highlight the importance of neuro-immune communication for the success and survival of all species. Apart from homeostatic functions, neuro-immune interactions are vital for the protection of neuronal tissues from invading pathogens as well as from damaging host immune responses. For instance, *Caenorhabditis elegans* responds to microorganisms by utilizing its nervous system, which triggers a protective behavioral avoidance response (Bargmann et al., 1990; Zhang et al., 2005). This behavior depends on G protein-coupled receptors (GPCRs) expressed by chemosensory neurons (McMullan et al., 2012).

Viral pathogens have evolved different strategies to invade the CNS including exploiting the olfactory route or crossing the BBB (Koyuncu et al., 2013; Mori et al., 2005). HSV-1, influenza A virus, parainfluenza viruses, are some examples of viral pathogens that enter the CNS through olfactory organ in mammals (Detje et al., 2009; Koyuncu et al., 2013; Mori et al., 2005). CNS immune responses need to be quick and tightly controlled because, otherwise, they may lead to meningitis, encephalitis, meningoencephalitis, or even death (Koyuncu et al., 2013). The present study reveals a novel model of viral recognition by the vertebrate nervous system in which OSNs are able to sense microbial-derived signals and use electrical decoding to trigger ultra-fast immune responses in teleosts.

We have identified a specific type of OSN, the crypt neuron, that acts as a canary in the coal-mine in the bony fish olfactory organ. We envisage 3 features of crypt neurons that make them ideal for pathogen detection: (i) they are strategically located in the most apical part of the teleost olfactory epithelium and so are the most exposed to invading microorganisms; (ii) they only constitute ~1-2% of all cells in the OO of trout, making TrkA-mediated cell death an exquisitely specific mechanism of triggering immune responses without compromising large numbers of OSNs; and (iii) because crypt neurons are replaced within few days, the refractory period to recover pathogen sensors in the trout OO is relatively quick. The effects of a secondary pathogen encounter prior to crypt neuron regeneration remain to be explored but it is possible that compensatory mechanisms are in place while crypt neurons replenish.

Viral-induced cell death allows crypt neurons to efficiently orchestrate innate immune responses such as induction of CCL19-like and the recruitment of CD8^+^ T cells to the local nasal environment. CD8^+^ T cells have been shown to prevent HSV-1 reactivation without destroying the infected sensory neurons in the trigeminal ganglia of mice (Liu et al., 2000). Thus, the CD8^+^ T cell infiltrates detected in the teleost OO after nasal IHNV delivery may play a critical role in the elimination of the virus without destruction of OSNs. Future studies will aim to further understand the role of fast-recruited CD8 T cells in the nasal mucosa.

TrkA molecular receptor is known to be utilized by several pathogens including parasites such as Trypanosoma (de Melo-Jorge and PereiraPerrin, 2007) and viruses such as HSV (Cabrera et al., 2015) in order to invade host neurons or to change neuronal behavior. Interestingly, secreted HSV G protein is able to bind TrkA in mouse skin neurons and modify neuronal dendrite outgrowth (Cabrera et al., 2015). Additionally, HSV-2 secreted G protein, is known to bind chemokines and enhance in this manner cell migration (Martínez-Martín et al., 2016). We provided evidence that the G protein of aquatic rhabdoviruses such as IHNV also interacts with TrkA^+^ crypt neurons. These findings suggest that G proteins from different neurotropic viruses have co-opted binding TrkA expressed in different neuronal types as a strategy to invade their hosts. Importantly, our work shows that the arms race between host and pathogen has resulted in efficient immune responses evoked by the TrkA-viral G protein interaction.

One of the most striking findings of the present study was the immunological cross-talk between the OO and the OB. Previous studies in mice have shown that infiltration of CCR7^+^ CD8^+^ T cells from the lymph nodes into the OB occurs in response to neurotropic viral infection of the OB (Cupovic et al., 2016). Our experiments revealed that the expression of CK10, a CCL19-like chemokine in trout (Sepahi et al., 2017), is quickly up-regulated on in the OO and down-regulated in the OB in response to nasal viral delivery and that zebrafish larvae where crypt neurons had been ablated showed dysregulated CCL19-like expression patterns. These findings suggest that the CCL19-CCR7 CD8 T cell axis is a conserved hallmark of viral neuronal infections in both bony fish and mammals. Importantly, the present study shows that a type of OSN, the crypt neuron, regulates antiviral immunity not only at the site of antigen encounter, but also in the OB, even in the absence of viral antigens in this tissue. Through a mechanism not explored in the present study, the OB turns viral-evoked electrical signals into immune responses within minutes. Future studies should address which molecules (i.e neurotransmitters or neuropeptides) are responsible for the changes in the OB microvasculature and T cell migration following nasal viral delivery.

In conclusion, our findings demonstrate a new mechanism of neuroimmune interaction by which OSNs can rapidly initiate antiviral immune responses in the OO-OB axis via a TrkA-sensing mechanism of viral G proteins. Understanding how the interaction between viral antigens and OSNs regulates innate and adaptive immune responses in the nasal mucosa and CNS can potentially help improve the efficacy and safety of nasal vaccines.

## Acknowledgements

The authors would like to thank Dr. E. Casadei for cloning the IHNV G protein, Hossein Goudarzi for his help with graphical abstract and I. Fuentes and P. Martínez for their excellent technical assistance, Dr. Pilar Fernández-Somalo for the SVCV strain, and Profs. D. Halpern and Parsons for the the Tg(UAS-E1b:nfsb-mCherry)^c264^, Dr. Weiming Li for his help with the preliminary data in the electrophysiology assays.

This work was supported by USDA AFRI Grant 2DN70-2RDN7 to IS, NSF IOS award 1755348 to IS, and NIH COBRE grant P20GM103452 as well as Spanish Ministry of Economy and Competiveness (grant BIO2014-52655-R and BIO2017-84702-R to VM), co-funded with Fondos Europeos de Desarrollo Regional/European Regional Development Funds. Cecelia Kelly was funded by the Stephanie Ruby Fellowship and NSF award 1456940 to IS.

## Author contributions

AS performed most of the trout experiments, analyzed data and wrote the manuscript, AK performed confocal microscopy, sequence alignments and zebrafish gene expression experiments, PM performed TEM and light microscopy experiments, JGV performed zebrafish infections and ablation experiments, CK and DGM made transgenic zebrafish, CJ made recombinant protein, MH performed electrophysiology experiments, VM conceived zebrafish experiments and wrote the manuscript, IS conceived the study, analyzed data and wrote the manuscript. All authors approved the manuscript before submission.

## METHODS

### Animals, nasal delivery of virus and tissue sampling

All rainbow trout studies were reviewed and approved by the Institutional Animal Care and Use Committee (IACUC) at the University of New Mexico, protocol number 16-200384-MC. For nasal delivery of virus studies, rainbow trout (mean weight of 50-150 g) received 30 μl of live attenuated infectious hematopoietic necrosis virus (IHNV) (2 x 10^8^ PFU/ml) or phosphate buffer saline (PBS) in each nare. For TrkA blocking experiment, rainbow trout (N = 20) received 30 μl of 10 μM AG879 or vehicle 30 min before viral delivery as described in diagram (Figure 1K). Olfactory Organ (OO) and olfactory bulbs (OB) were snap frozen and cryoblocks used for immunostaining or kept in RNAlater for gene expression studies.

Zebrafish (*Danio rerio* H.) were obtained from the Zebrafish International Resource Center and mated, staged, raised and processed as described (Westerfield, 2000). The line *Tg(UAS-E1b:nfsb-mCherry)^c264^* was previously reported (Davison et al., 2007) The experiments performed comply with the Guidelines of the European Union Council (Directive 2010/63/EU) and the Spanish RD 53/2013. Experiments and procedures were performed as approved by the Bioethical Committees of the University of Murcia (approval numbers #537/2011, #75/2014 and #216/2014).

### Electrophysiological recordings

Rainbow trout were anesthetized in a solution of MS222 at 0.1 g/l, and then immobilized with an intra-muscular injection of gallamine triethiodide (3 mg/kg of body weight, in 0.9% saline). Fish were then secured in a V-shape Plexiglas stand partially inundated, whereby gills could be continuously irrigated with aeriated anesthetic solution of MS222 at 0.05 g/l. The olfactory rosette was surgically exposed and borosilicate electrodes, filled with a solution of 3 M KCL in

0.4% agar and connected to solid state electrodes with Ag/AgCl pellets, were placed between olfactory lamellae (signal electrode) and external skin (reference electrode). The olfactory epithelium was continuously irrigated with tap charcoal filtered water and the stimulus was released directly into the nose through a borosilicate tube. The olfactory responses generated after release of the stimuli for 4 s were filtered and amplified by a NeuroLog DC filter and pre-amplifier integrated by an Axon Digidata 1550B, and stored on a PC running Axoscope 10.6 software.

#### Dose response experiments

Stimuli were serially diluted from a 1:100 to 1:1000 000 from a stock solution, and applied to the nose to measure amplitude of the olfactory responses. These responses were blank subtracted (i.e. the response to tap charcoal filtered water) and normalized to those of _*L*_-serine at 10^−5^ M. IHNV stock was a vaccine solution with inactivated IHNV at 2 x 10^8^ PFU and culture medium stock was the supernatant of the vaccine after being centrifuged at 50000 rpm for 20 min at 5 °C.

#### Cross-adaptation experiments

We identified dilutions of IHNV and medium that evoked the same EOG amplitude (called the ‘unadapted’ response). Then the olfactory rosette was continually exposed to IHNV solution at the concentration of the unadapted response at least 1 min, and the response to a sample at double concentration of unadapted response IHNV was recorded (called the self-adapted control, SAC). After that, the response to a mixture IHNV and medium, both at same concentration that unadapted response, was recorded (Mix). Both measures, Mix and SAC, were then calculated as a percentage of the unadapted response. After adaptation, the olfactory rosette was flushed with charcoal filtered water for 20 min, and the process repeated using medium as the adapting solution and IHNV or the mixture IHNV and medium as stimuli. Half of the fish were adapted first to IHNV and the other half first to control.

#### Inhibition curves

Responses to 1:1000 to IHNV or medium were recorded (both showed similar amplitude in their olfactory responses). Then the olfactory rosette was continuously exposed to increasing concentrations of AG879 from 10^−9^ M to 10^−5^ M and, under each adapting concentration of the drug, it was measured the olfactory responses to 1:1000 of IHNV or medium. Responses were calculated as ratio between 1:1000 odorant after adaptation to drug solution and 1:1000 odorant before adapted to drug. All graphs were produced with Sigma plot 11.0 and EC50 concentrations were calculated using the Pharmacology module of the same program.

### Histology, transmission electron microscopy and immunofluorescence microscopy

For transmission electron microscopy (TEM), the OO (N = 3) of rainbow trout that had received live attenuated IHNV IN 15 min prior to sampling were fixed overnight at 4 °C in 2.5 % (v/v) glutaraldehyde in PBS, then transferred to 1 % osmium tetroxide (w/v) in PBS for 2 h at 4 °C. After washing in PBS (3 times, 10 min), samples were dehydrated in a graded series of ethanol (10–100 %) through changes of propylene oxide. Samples were then embedded in Epon resin, sectioned and stained with uranyl acetate and lead citrate before being examined in a PHILIPS TECNAI 12 transmission-electron microscope. Additionally, semithin sections were stained with toluidine blue. The conjugated antibodies used for immunostaining are listed in the **Key Resources Table**. Trout OO and OB were snap frozen in OCT and 5-μm-thick cryosections were fixed in 4% paraformaldehyde for 3 min, blocked and labeled with rat anti-trout CD8α (1:50 dilution), anti-pERK (1:50 dilution), anti-human TrkA (1:100 dilution) and rabbit anti-mouse caspase 3 (1:100 dilution) antibody for immunostaining. Nuclei were stained with DAPI. Samples were observed under a Nikon Ti or Zeiss confocal microscopes. To test the permeability of BBB 15 min after nasal viral delivery, 50 μl of FITC-conjugated 10 kDa dextran particles in PBS were injected i.v into 10 g rainbow trout (N = 6) 1 hour before sampling. Trout then received IHNV or PBS IN and 15 min later, trout heads were snap frozen, embedded in OCT and cryosections were examined for fluorescence microscopy.

### Western blot, cell isolation and flow cytometry

The conjugated antibodies used for western blot and Flow cytometry are listed in the **Key Resources Table**. Olfactory Organ (OO), head kidney (HK), and brain (B) were extracted and prepared for Western blotting as explained elsewhere (Sepahi et al., 2016b). Briefly, tissues were lysed in RIPA buffer and 10 μl of lysed tissues were mixed with 10 μl of Laemmli buffer under non-reducing conditions. Samples were boiled for 3 min at 97 °C and resolved on 4–15% SDS-PAGE gels. Gels were run for 50 min at 120 V and transferred onto PVDF membranes. Membranes were blocked in PBS-T containing 5% non-fat milk overnight at 4 °C. Membranes were incubated with anti-TrkA (1:1000) for 90 min, washed three times in PBS-T and then incubated for 60 min with HRP-anti-rabbit IgG (1:2500). Detection was performed using ECL Western Blotting Substrate. Immunoblots were scanned using a ChemiDoc Touch Imaging System and band densitometry was analyzed with Image Lab Software.

Isolation of trout OO cells was carried out as explained elsewhere (Tacchi et al., 2014). Briefly, trout OO were obtained by means of mechanical agitation of both olfactory rosettes in DMEM medium (supplemented with 5% FBS, 100 U/ml penicillin and 100 μg/ml streptomycin) at 4 °C for 30 min. Leukocytes were collected, and the aforementioned procedure was repeated four times. Thereafter, the OO pieces were treated with PBS (containing 0.37 mg/ml EDTA and 0.14 mg/ml dithiothreitol) for 30 min followed by enzymatic digestion with collagenase (0.15 mg/ml) for 2 h at 20 °C. All cell fractions obtained the OO following the mechanical and enzymatic treatments were pooled, washed with modified DMEM. OB microvessels were extracted then 40μl of DMEM containing heparin were added to the cavity to collect the remaining blood from the microvessels. This step was repeated twice. Cells were isolated by forcing the tissue through a 100-μm pore nylon cell strainer and washed in DMEM three times. Cell suspensions were counted in a haemacytometer and stained with CD8α, IgM and IgT antibodies as explained elsewhere (Sepahi et al., 2016a). After washing, a total of 30,000 cells were recorded using an Attune NxT flow cytometer. The percentage of CD8α^+^, IgM^+^ and IgT^+^ cells was quantified as the percentage of FITC^+^ cells within the lymphocyte gate using their FSC/SSC profile.

### Gene expression analysis by real-time quantitative PCR

Total RNA from OO and OB samples were collected and placed in 1 ml TRIzol for RNA extraction according to the manufacturer’s instructions. cDNA synthesis was performed as explained elsewhere (Sepahi et al., 2016b). The resultant cDNA was stored at -20 °C. The expression of *pgs2b, IFN-γ*, TNFα*, CK10* (CCL19-like) and *c-fos* for rainbow trout and *CCL19-like* and SVCV N protein for zebrafish were measured by RT-qPCR using specific primers (**Key Resources Table**). The qPCR was performed using 3 μl of a diluted cDNA template as described in (Tacchi et al., 2013). The relative expression level of the genes was determined using the Pfaffl method (Pfaffl, 2001)as previously described (Tacchi et al., 2013).

### DNA constructs and generation of transgenic zebrafish larvae

Three kb of the upstream regulatory sequence of the *ora4* gene, which includes its 5’UTR (Ensembl accession number ENSDARG00000078223) was amplified using PfuUltra II Fusion HS DNA Polymerase, the primers 5’-aaggtaccgtgaatgcgtgtgtgtgatgtc-3 and 5’-aaaggatccgctgaagatgctccagagtcc-3, and zebrafish larval genomic DNA as template. The amplicon was digested with *KpnI*/*BamHI*, cloned in the p5E-MCS vector (#228) of the Tol2kit and the *ora4::Gal4VP16* construct were then generated by MultiSite Gateway assemblies using LR Clonase II Plus according to standard protocols and using Tol2kit vectors described previously (Kwan et al., 2007).

The lines *Tg(ora4::Gal4VP16)^ums4^* were generated by microinjecting 0.5-1 nl into the yolk sac of one-cell-stage embryos a solution containing 100 ng/μl *ora4::Gal4VP16* construct and 50 ng/μl Tol2 RNA in microinjection buffer (0.5x Tango buffer and 0.05 % phenol red solution) using a pneumatic microinjector.

### Cell ablation and live imaging of zebrafish larvae

The lines *Tg(UAS-E1b:nfsb-mCherry)^c264^* and *Tg(ora4::Gal4VP16)^ums4^* were crossed. Their offspring were treated at 48 h post-fertilization (hpf) for 24 h with 12 mM metronidazole and kept in dark (Davison et al., 2007). Images were first obtained at 48 hpf as described below. Thereafter, starting from 72 hpf the prodrug was removed, and larvae imaged once a day up to 120 hpf to confirm the ablation of ora4^+^ crypt neurons.

For live imaging, larvae were anesthetized in tricaine as previously described (Galindo-Villegas et al., 2012). Images were captured with an epifluorescence MZ16FA stereomicroscope (Leica) equipped with green and red fluorescent filters while animals were kept constantly at 28.5°C.

### Viral challenge in zebrafish

The SVCV isolate 56/70 was propagated in EPC cells and titrated in 96-well plates. Thirty 72 hpf zebrafish larvae per group in triplicate were challenged for 24 h at 25°C in disposable Petri dishes by immersion in 10^8^ TCID50/fish SVCV. After challenge, the remaining fish in each group were transferred to fresh plates containing egg water and monitored every 12 h over a 6-day period to score mortality (López-Muñoz et al., 2010).

### Statistical analysis

Results are expressed as the mean ± SE. Data analysis was performed in GraphPad Prism version 5.0. The RT-qPCR measurements were analyzed by *t*-test to identify statistically significant differences between groups. One-way ANOVA and a Tukey post hoc analysis test were performed to identify statistically significant differences among groups. Statistical analysis for survival assay was carried out using PRISM 7 for Mac OS X (GraphPad). Gehan-Breslow-Wilcoxon method was performed following a log-rank test and confirmed with Kaplan-Meier curve to ensure compatibility and avoid deviations due a lack of proportional hazard. P-value of < 0.05 was considered statistically significant.

### Contact for reagent and resource sharing

Further information and requests for resources and reagents should be directed to and will be fulfilled by the Lead Contact, Irene Salinas

---

Supplementary Fig. 1: (A) Detection of TrkA in trout OO and brain but not HK lysates by immunoblotting. Immunoblots detecting TrkA showed a band at the expected size (~140 KDa) in OO and brain. (B) Immunofluorescence staining of control rainbow trout HK cryosection stained with anti-TrkA antibody (FITC, green) confirming absence of TrkA^+^ cells. Cell nuclei were stained with DAPI DNA stain (blue). Scale bar: 20 μm.

Supplementary Fig. 2: IHNV activates sensory neurons in the OO in vitro (A) Representative dot plots of control (left) and IHNV (right) trout OO extracted cells stained with anti-pERK antibody showing the mean percentage of positive cells. (B) Quantification of flow cytometry data in (A) indicating a significant increase in the percentage of pERK^+^ cells 15 min after adding IHNV (multiplicity of infection 1:3) *in vitro*. Results are representative of three independent experiments (N = 5). *p < 0.05

Supplementary Fig. 3: Nasal delivery of IHNV does not result in presence of virus in the OB and does not alter BBB integrity 15 min after delivery. (A) Immunofluorescence staining with anti-IHNV Abs (Cy3, red) showing no IHNV staining in the OO of control rainbow trout. (B) Immunofluorescence staining with anti-IHNV Abs (Cy3, red) showing the presence of IHNV (red arrows) in the OO of IHNV treated rainbow trout 15 min after nasal delivery. (C) Immunofluorescence staining with anti-IHNV Abs (Cy3, red) showing no detection of IHNV at OB of control rainbow trout. (D) Immunofluorescence staining with anti-IHNV Abs (Cy3, red) showing the absence of IHNV in the OB of IHNV treated rainbow trout 15 min after nasal delivery. Scale bar, 20 μm. (E-H) Intravenous injection of FITC-conjugated dextran showing that no changes in the BBB integrity in IHNV-treated fish as demonstrated by the absence of FITC staining in the OB. Scale bar: 100 μm.

Supplementary Fig. 4: Leukocyte recruitment occurs as early as 15 min after IHNV delivery as visualized by enlargement lamina propria (LP) of the olfactory lamellae of IHNV-treated compared to control fish. (A) Immunofluorescence staining of control (left) and IHNV-treated (right) rainbow trout OO stained with anti-trout TrkA (FITC, green) showing our image analysis strategy and the enlargement in the apical and medial regions of the LP in the IHNV-treated fish. Cell nuclei were stained with DAPI DNA stain (blue). Results are representative of two different experiments (N = 3). Scale bar, 20 μm. (B) The width of LP at the apical (100 μm from the lamellar tip) and lateral (250 μm from the lamellar tip) regions of the olfactory lamella were measured by image analysis of 10 individual lamellae from three different fish per treatment. The mean distance ± SE is shown. (C) Representative hematoxylin-eosin stain of adult rainbow trout olfactory organ showing Leukocyte recruitment occurs as early as 15 min after IHNV delivery of the olfactory lamellae of IHNV-treated (middle and right) compared to control fish (left). L, lumen; LP, lamina propria. Scale bar: 50 μm.

Supplementary Fig. 5: A low degree of amino acid conservation between IHNV G protein and HSV secreted G protein indicated. (A) Amino acid sequence alignment of HSV-2 sG protein (accession number GD_HHV23) and trout IHNV G protein (sequenced obtained from the live attenuated IHNV used in this study) performed in CLUSTALW showing a low degree of amino acid conservation (B) Production of recombinant FLAG-tagged IHNV G protein by mammalian expression system. Immunoblot using anti-Flag antibody confirmed the presence of the recombinant protein (IHNV G protein) band at expected (~50 KDa) molecular weight.

